# Isogenic hiPSC models of Turner syndrome development reveal shared roles of inactive X and Y in the human cranial neural crest network

**DOI:** 10.1101/2023.03.08.531747

**Authors:** Darcy T. Ahern, Prakhar Bansal, Isaac V. Faustino, Heather R. Glatt-Deeley, Rachael Massey, Yuvabharath Kondaveeti, Erin C. Banda, Stefan F. Pinter

## Abstract

**SUMMARY:** Modeling the developmental etiology of viable human aneuploidy can be challenging in rodents due to syntenic boundaries, or primate-specific biology. In humans, monosomy-X (45,X) causes Turner syndrome (TS), altering craniofacial, skeletal, endocrine, and cardiovascular development, which in contrast remain unaffected in 39,X-mice. To learn how human monosomy-X may impact early embryonic development, we turned to human 45,X and isogenic euploid induced pluripotent stem cells (hiPSCs) from male and female mosaic donors. Because neural crest (NC) derived cell types are hypothesized to underpin craniofacial and cardiovascular changes in TS, we performed a highly-powered differential expression study on hiPSC-derived anterior neural crest cells (NCCs). Across three independent isogenic panels, 45,X NCCs show impaired acquisition of PAX7^+^SOX10^+^ markers, and disrupted expression of other NCC-specific genes, relative to their isogenic euploid controls. In particular, 45,X NCCs increase cholesterol biosynthesis genes while reducing transcripts that feature 5’ terminal oligopyrimidine (TOP) motifs, including those of ribosomal protein and nuclear-encoded mitochondrial genes. Such metabolic pathways are also over-represented in weighted co-expression gene modules that are preserved in monogenic neurocristopathy. Importantly, these gene modules are also significantly enriched in 28% of all TS-associated terms of the human phenotype ontology. Our analysis identifies specific sex-linked genes that are expressed from two copies in euploid males and females alike and qualify as candidate haploinsufficient drivers of TS phenotypes in NC-derived lineages. This study demonstrates that isogenic hiPSC-derived NCC panels representing monosomy-X can serve as a powerful model of early NC development in TS and inform new hypotheses towards its etiology.

## INTRODUCTION

The absence of the second sex chromosome (monosomy-X) represents the only viable human monosomy in humans, resulting in Turner syndrome (TS, 1:2000 live births). Although fewer than 0.3% of conceptuses with monosomy-X (45,X karyotype) complete development ^1,2^, it is considered viable, because most sex-linked genes are expressed from a single copy in placental mammals: the Y lost most of its ancestral genes originally shared with the proto-X, except for dosage-sensitive regulators ^3^, while the X became subject to dosage-compensation in the form of X chromosome inactivation (XCI) in XX females ^4^. However, many genes retained on the Y have a homologous X-linked copy that escapees XCI. In addition to these X/Y-pairs (“gametologs”), genes in the recombining pseudoautosomal region (PAR) shared by X and Y also escape XCI in XX females ^4^.

The compound haploinsufficiency of these gametolog and PAR genes is hypothesized to underpin both cardinal short stature and premature ovarian failure in persons with TS, alongside variably penetrant (40-80%) phenotypes ^5^, like lymphedema of hands and feet, neurocognitive changes, as well as renal and cardiac malformations. Joining skeletal and craniofacial changes, bicuspid aortic valve and aortic defects in TS implicate cell types derived from neural crest cells (NCCs). These transitory multipotent progenitors arise from the dorsal side of the closing neural tube and migrate ventrally to contribute to the developing face, skin, heart, adrenal glands, joints and peripheral nervous system ^6^. Because all of these organ systems are variably impacted in subsets of TS patients ^7^, the neural crest is thought to be particularly vulnerable to monosomy-X.

Yet, direct experimental evidence for this hypothesis has been lacking. Because many murine orthologs of human PAR genes reside on autosomes ^8,9^, their gene dosage is maintained in X-monosomic mice, which consequently fail to model most TS phenotypes ^10,11^. To address this problem, we derived and cytogenomically validated X-monosomic human induced pluripotent stem cells (hiPSCs) alongside isogenic euploid control lines from mosaic fibroblasts. We recently reported that such isogenic panels from three unrelated donors recapitulate a hypothesized ^2^ impairment of trophoblast cell fates due to monosomy-X ^12^.

Here, we derive anterior NCCs from each of these hiPSC panels to assess the impact of human monosomy-X on the neural crest. We perform an in-depth transcriptomic and cellular analysis comparing monosomy-X to their otherwise isogenic euploid controls, and find monosomy-X hiPSCs give rise to significantly fewer PAX7/SOX10 double-positive NCCs compared to 46,XY euploid controls. Differential expression reflecting monosomy-X is highly concordant across male and female-derived panels, and jointly enriched in pathways associated with cholesterol metabolism, NCC-relevant cell signaling, translation and mitochondrial function. Importantly, NCC gene modules that reflect monosomy-X by weighted gene co-expression network analysis (WGCNA) are also preserved in hiPSC-derived NCC models of monogenic conditions traditionally considered to represent neurocristopathy.

## RESULTS

Over half a century of detailed studies in the chick and other model systems have shed light on the signaling pathways and intricate gene regulatory networks (GRNs) that govern the specification and migration of NCCs ^13,14^. These exquisite developmental studies have also informed progressively intricate protocols for NCC derivation from embryonic stem cells and hiPSCs that recapitulate terminal NCC fates as a function of their axial identity ^15–17^. Due to our interest in the anterior/craniofacial and vagal neural crest, which populate the developing face and heart, we applied a highly robust method for NCC specification via moderate WNT and BMP activation that is reported to enable reproducible derivation of anterior NCCs at ∼ 70% efficiency ^17,18^. We applied this method to a total of 11 distinct hiPSC lines from all three of our cytogenomically validated isogenic hiPSC panels ^12^, and quantified the dorsal neural tube marker PAX7 alongside the canonical neural crest marker SOX10, which jointly mark NCC specification. Applying automated immunofluorescence quantification, we observe the expected rate of PAX7^+^ SOX10^+^ double-positive cells (60-70%) across all euploid lines, though female euploid lines with somewhat reduced efficiency relative to male euploid lines (Fig. 1A). While monosomy-X hiPSCs can also differentiate to PAX7^+^SOX10^+^ NCCs, their overall efficiency is significantly reduced by about half in both male-derived isogenic panels (Fig. 1B, male 1: p=0.002, male2: p=1.1×10^-5^, Mann-Whitney U). Interestingly, female-derived monosomy-X NCCs arise at a similar rate as their euploid counterparts, rendering this isogenic comparison non-significant (p=0.3) but still significantly reduced relative to male euploid controls (p≤1.3×10^-3^). Specifically, we find the PAX7^+^ rate is largely unaffected by monosomy-X across all three panels (Fig. S1), whereas it is SOX10^+^ in male1 and male2-derived monosomy-X lines that is significantly reduced (male 1: p=0.001, male2: p=9.8 ×10^-6^, Mann-Whitney U).

**Figure 1:**
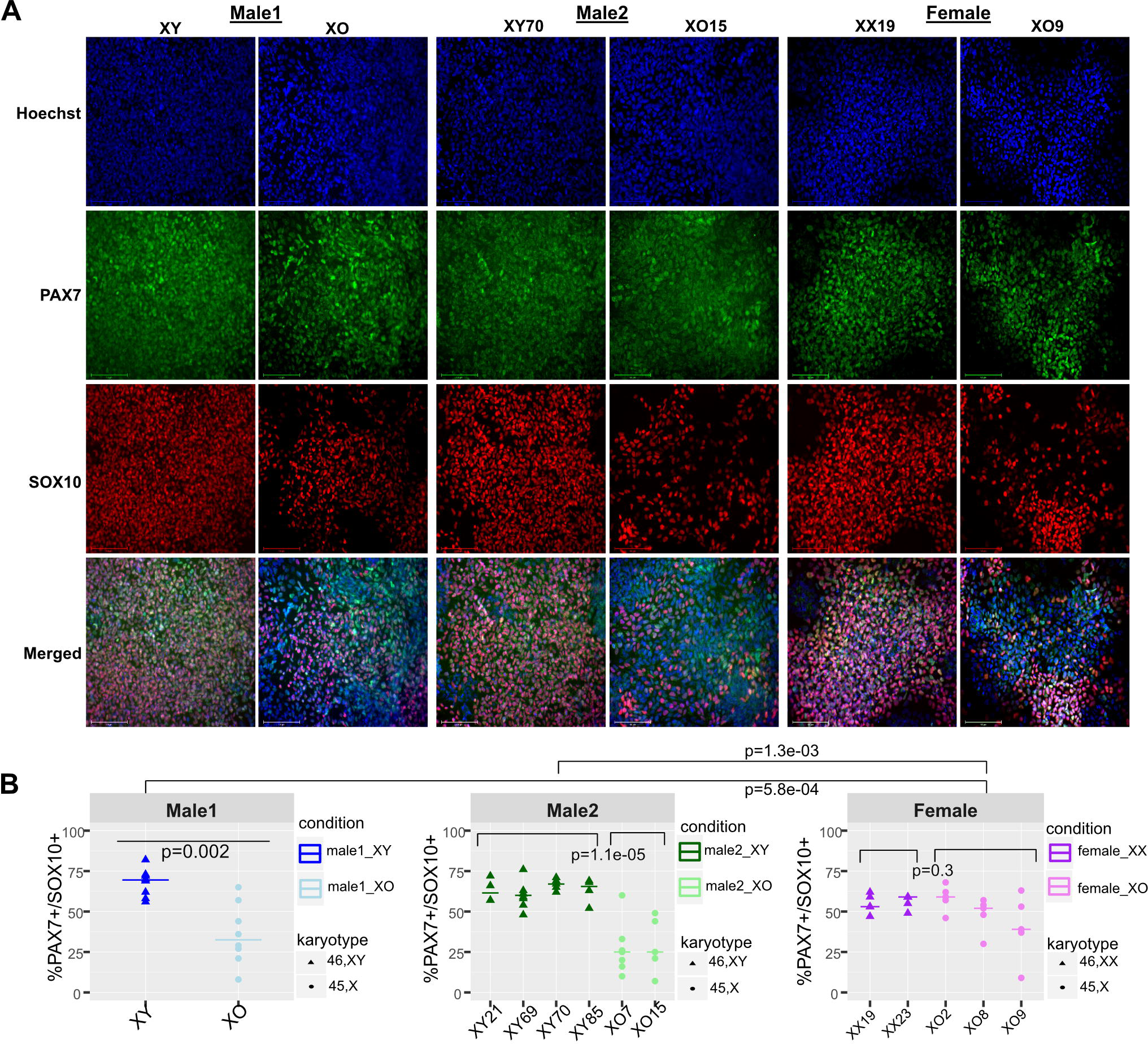
Monosomy-X impact on neural crest differentiation relative to isogenic euploid controls. **(A)** Representative immunofluorescence (IF) of PAX7 (green), SOX10 (red) and nuclei (Hoechst33342) in hiPSC-derived pairs of 45,X and euploid control NCCs from Male1/2 and Female donors (100 um scale bar). **(B)** CellProfiler quantification of 4-7 rounds of differentiation. Brackets and p-value (Mann-Whitney U test) indicate grouped cell lines compared within and across isogenic panels (colored by donor, symbols denoting karyotype).

We next performed a highly-powered RNA-seq study on a subset of these samples, with a minimum of 4 replicates per cell line, grouped by a combination of donor and karyotype (“condition”), comprising 7-12 replicates each. This enables our analysis to assess monosomy-X driven changes within each isogenic context (or donor). Indeed, each paired set of 45,X and euploid controls segregate from each other along the first and second principal components (Fig. 2A), while hierarchical clustering groups all male-derived monosomy-X samples together (Fig. S2A). We next compared X-monosomic and euploid NCC expression levels of marker gene sets identified in two prior human NCC differentiation studies ^15,18^, orthologs of chick neural crest GRNs ^19^, and genes differentially expressed (DEGs) in the murine neural crest ^20^, as well DEGs of a 3D *in vitro* model of the folding human neural tube ^21^. Indeed, gene sets from these five studies were largely sufficient to segregate euploid and X-monosomic samples (Fig. S2B). Single-cell DEGs of the human 3D neural tube model were the most highly expressed, while early anterior neural crest (eANC), murine Hox-negative NC markers, and migratory NC markers identified in the chick were roughly tied. We also observed uniformly low expression of posterior HOX genes, confirming the anterior identity of our NCCs (Fig. S2C). Assessing median-normalized gene set levels, we observe significantly lower levels of early anterior neural crest (eANC) and higher levels of late ANC (lANC) markers in euploid over X-monosomic lines in all three isogenic panels (Fig. 2B), and respectively higher expression of NC over neuroectodermal genes (NE) in male euploid over X-monosomic NCCs (Fig. S2D). Likewise, genes expressed in p75^+^ hESC-derived NCCs are also significantly higher in euploid over X-monosomic hiPSC-derived NCCs, which is also true for orthologs of GRNs identified in the chick migratory neural crest (Fig. 2C,D).

**Figure 2:**
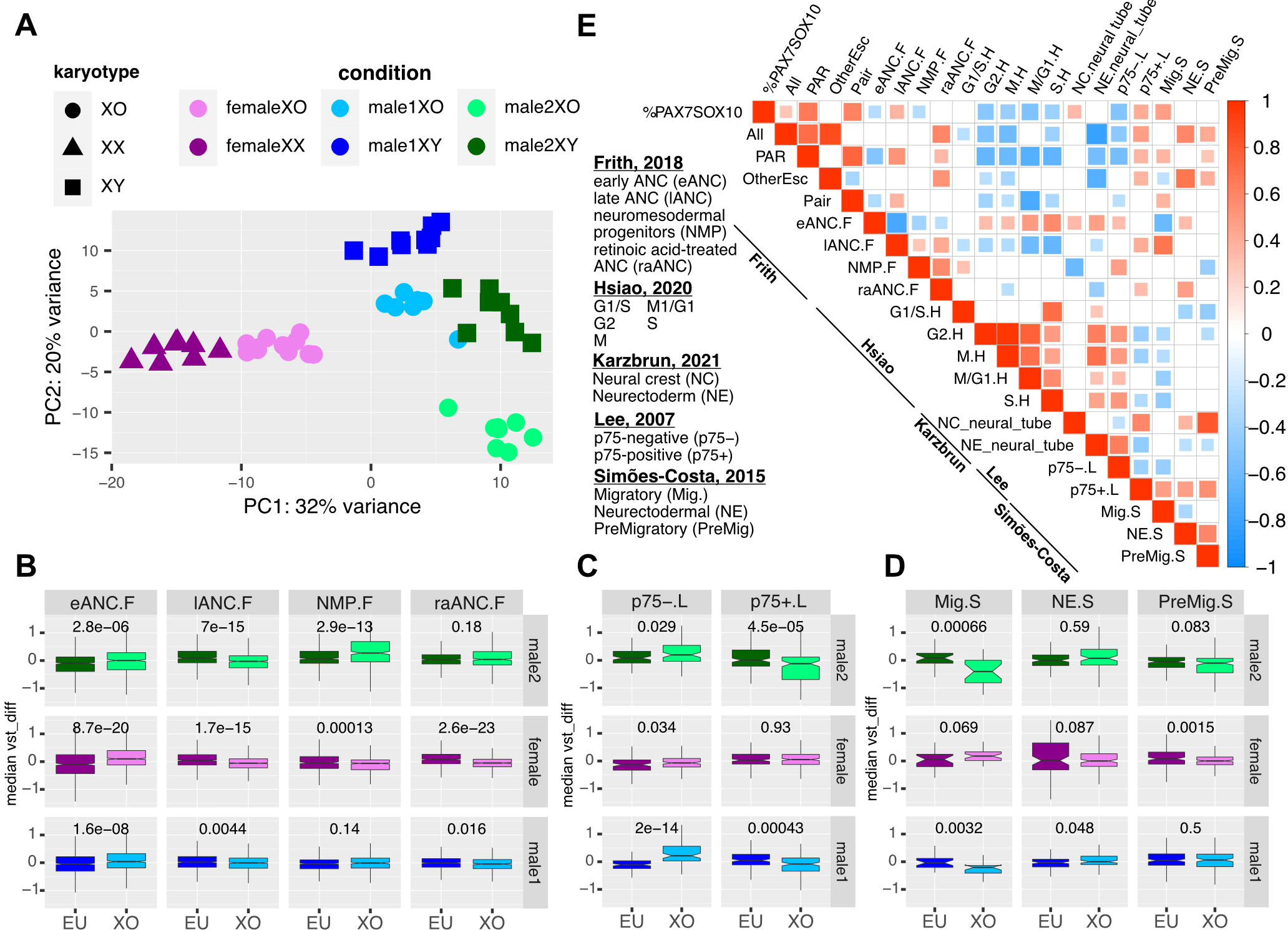
Reduction of neural crest marker expression in monosomy-X relative isogenic euploid controls. **(A)** Principal component analysis (PCA) segregates samples by “condition” (donor & karyotype), with % of variance of PC1/2 as indicated. **(B)** Variance-stabilized counts (vst) of early and late anterior NC (e/lANC), retinoic acid-treated NC (raANC) and neuromesodermal progenitor (NMP) markers from ^18^ (vertical panels) were median-normalized and averaged over euploid (EU) and 45,X (XO) NCCs of each (horizontal) donor panel (significant differences denoted by Mann-Whitney P-value). **(C,D)** as in B) for respectively p75-/+ associated markers ^15^, and chick NC marker sets ^19^. **(E)** Pearson correlation of the matching PAX7^+^SOX10^+^ percentage with averaged marker sets from (B-D), alongside cycling markers ^59^, NC markers from a 3D folding human neural tube model ^21^, and averaged pseudo-autosomal, X/Y Pair, and other escapee (PAR/PAIR/oESC) expression. Only significant pairwise correlation shown (p≤0.05, Fisher transformed Pearson R).

To delineate which groups of X/Y-linked genes may be haploinsufficient in NCCs, we plotted PAR and gametolog pair genes alongside X-specific genes that escaped XCI in female euploid NCCs, as reflected in allelic variant counts of the phased female X (Fig. S2E). Largely driven by PAR genes, monosomy-X NCCs segregate from their isogenic euploid controls, which expectedly split female 46,XX and male 46,XY samples. DEGs escaping XCI in euploid female NCCs (lesser allele frequency, LAF≥0.1, binomial p≤0.05) included 7 PAR1 genes, 5 X-linked gametologs and 16 previously-reported escapee genes, alongside 8 genes that may reflect novel NCC-specific escapees or have reactivated in a subset of cells despite robust bulk expression of *XIST* (Fig. S2F). In male 46,XY hiPSCs, Y-linked gametologs generally trailed their X-linked homolog in expression by a limited (∼2-3) vst differential (Fig. S2G), with some nearly-equal (*NLGNX/Y & TXLNG/Y*) and more divergent exceptions (*TMSB4X/Y, TBL1X/Y*). We next correlated the PAX7^+^ SOX10^+^ fraction with the median expression of PAR, Pair, other escapee genes and their summed expression (“All”) in each sample (Fig. 2E). Including the NCC marker gene sets in this analysis, we observe that the PAX7^+^ SOX10^+^ fraction best correlates with levels of PAR and Pair genes, which likewise correlate with the late ANC, p75^+^ and chick migratory GRN sets. Because NCC undergo an epithelial-to-mesenchymal transition that modulates cell cycle progression ^22,23^, more mature NCC markers are expectedly anti-correlated with cell cycle and earlier NCC markers. In sum, these data suggest monosomy-X hinders or delays the maturation of early to late neural crest, in keeping with their reduction of SOX10^+^ positive cells.

To determine whether global differential expression points to a common monosomy-X signature across isogenic panels, we called and compared DEGs common to any two or all three panels. Altogether, over 30% of all DEGs were shared in all pairwise comparisons (p = 1.1×10^-13^, p = 1.2×10^-21^ and p = 3.7×10^-137^, hypergeometric test), with the two male-derived monosomy-X panels reflecting the most significant overlap (Fig. 3A). We also assessed the direction of gene expression changes in these overlapping DEG sets, which perfectly segregate all monosomy-X samples from their isogenic euploid controls. Indeed, DEGs shared by all three (p = 3.7×10^-124^, sign test), as well as any two isogenic panels (p = 7.4×10^-195^, 5.1×10^-11^, 8.0×10^-47^) change concordantly in highly significant fashion (Fig. 3A). This level of overlap and concordant change is remarkable given that monosomy-X samples (‘mXO1/2’ & ‘fXO’) were only assessed relative to euploid controls in their own isogenic context.

**Figure 3:**
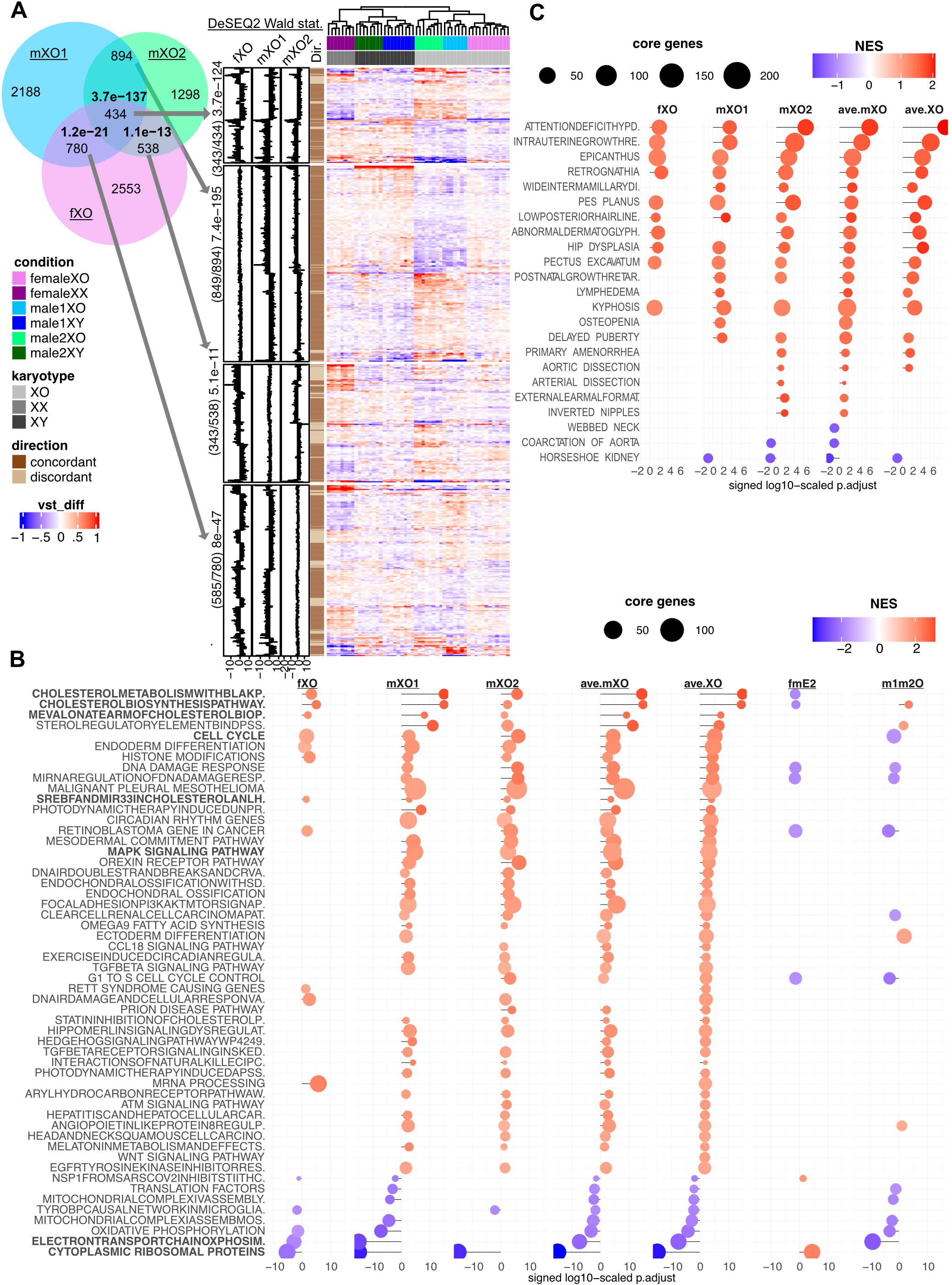
Concordant impact of monosomy-X on NCC transcriptomes across isogenic panels. **(A)** *Left:* Venn diagram of differentially expressed genes (DEGs) in male1, male2 and female (mXO1, mXO2) 45,X NCCs. Significance of pairwise overlapping DEGs in bolded p-values (hypergeometric distribution). *Right:* Differential vst heatmaps for overlapping DEG sets from (A) as denoted by arrows. Ratio and p-value (sign-test) denote the number of DEGs with concordant direction (“Dir.”) in the triple and all pairwise overlapping DEG sets, also shown in divergent (fXO, mXO1/2) annotation panels showing the Wald statistic (DESeq2) for each gene. Dendrogram segregates samples by karyotype, irrespective of donor. **(B)** Wikipathway gene set enrichment analysis (GSEA), ordered by the quantile-normalized mean Wald statistic across 45,X conditions (ave.XO). X-axis denotes log-scaled GSEA adjusted p-value. Colored by normalized enrichment score of up-(red) and down-(blue) regulated gene sets (sized by number of genes). Results for all isogenic monosomy-X comparisons (fXO, mXO1/2), averaged (ave.XO/ave.mXO) and control comparisons (fmE1/2: female-male euploids, and m1m2O: male1/2 45,X samples). **(C)** Human Phenotype Ontology (HPO) GSEA results for significantly (p.adjust≤0.1) enriched terms associated with Turner Syndrome (Orphanet ID: 881), ordered by the quantile-normalized mean (mXO1/2) Wald statistic (ave.mXO, otherwise as in B).

To identify cellular pathways and developmental processes commonly impacted by the lack of X or Y, we performed gene-set enrichment analysis (GSEA) for all three monosomy-X NCC panels as ordered by their averaged Wald statistic (‘ave.XO’). We also compared female-to-male euploid (‘fmE1/2’), as well karyotypically-identical male1-to-male2 monosomy-X (‘m1m2O’) NCCs. Mirroring the global overlap of DEGs (Fig. 3A), significantly enriched gene-sets revealed a highly concordant pattern across all three isogenic panels (Fig. 3B). Across Wikipathway (p.adj<0.02), KEGG and Hallmark (Fig. S3, Table S1) gene term collections, monosomy-X NCCs upregulate genes associated with cell cycle, cholesterol and lipid biosynthesis, and down-regulate genes linked to translation and oxidative phosphorylation. While the impact of monosomy-X in the female NCC panel is generally milder than the two male panels, the concordance in GSEA terms mirrors the highly significant overlap in DEGs, and both hyperlipidemia and hypercholesteremia are also seen in patients with TS ^24^. Likewise, in the human phenotype ontology (HPO), monosomy-X NCCs show dysregulated gene sets relating to craniofacial, joint, and cognitive development that are also frequently observed in patients with Turner Syndrome (Table S1), including ‘Down-slated palpebral fissures’, ‘joint hypermobility, ‘hyperactivity’ (all p.adj<0.0001). Indeed, the enrichment of HPO terms relating to Turner Syndrome (“ORPHA881”) in monosomy-X NCC vs. control comparisons is particularly striking (Fig. 3C), and recovers 28/114 (24.6%) of all HPO terms linked to TS (Table S1), none of which were significant in any of the control comparisons. In sum, these GSEA results underscore that differential expression in X-monosomic NCCs relative to euploid controls recovers gene terms of known TS phenotypes that relate to the neural crest, as well as beyond, and implicates cellular and developmental pathways impacted by monosomy-X.

We next performed weighted gene co-expression network analysis (WGCNA) to elucidate the relationships between monosomy-X and altered pathways, and prioritize individual X/Y-linked genes as potential dosage-sensitive drivers. This analysis assigned genes to 29 modules, 23 of which are driven by contrasting expression of euploid and X-monosomic expression that we define as a loss of module preservation (Z-score drop ≥5) in networks lacking euploid samples (Fig. S4A). Eight modules (groupA: 1,6,7,13,19,20,26,27) are significantly correlated (R≥0.3, p≤0.05) with the PAX7^+^SOX10^+^ rate, and six modules (groupB: 3,5,8,9,11,15) are significantly anti-correlated (R≤–0.3, p≤0.05, Fig. 4A). Consistently, the PAX7^+^SOX10^+^ correlated groupA modules associate with mature NCC markers, whereas PAX7^+^SOX10^+^ anti-correlated groupB modules reflect an earlier, NE-biased cycling cell signature. The PAX7^+^SOX10^+^ positively-correlated groupA was enriched in translation and mitochondrial function terms, while the PAX7^+^SOX10^+^ anti-correlated groupB was over-represented for cholesterol biosynthesis, cell cycle-related terms, and several signaling pathways, including p53, MAPK and mTOR signaling (Fig. 4A, S4B, Table S2). To further contextualize the relationship between these modules and biological processes, we turned to a semantic similarity map of significantly-enriched GO terms (p.adjust≤0.01, hypergeometric distribution) in modules that also significantly co/anti-correlated with the PAX7^+^SOX10^+^ rate (Fig. 4B, -/+ labeled modules). Terminal neural-crest related terms (“cranial facial parasympathetic nerve”, “endocardial cushion”, “anterior/posterior pattern specification”, “angiogenesis”) were enriched in groupB modules, whereas metabolic functions that connect cell cycle and splicing to translation and mitochondrial function were represented by both group A&B modules. Overall, these enriched metabolic terms mirrored the GSEA results (Fig. 3B, Table S1).

**Figure 4:**
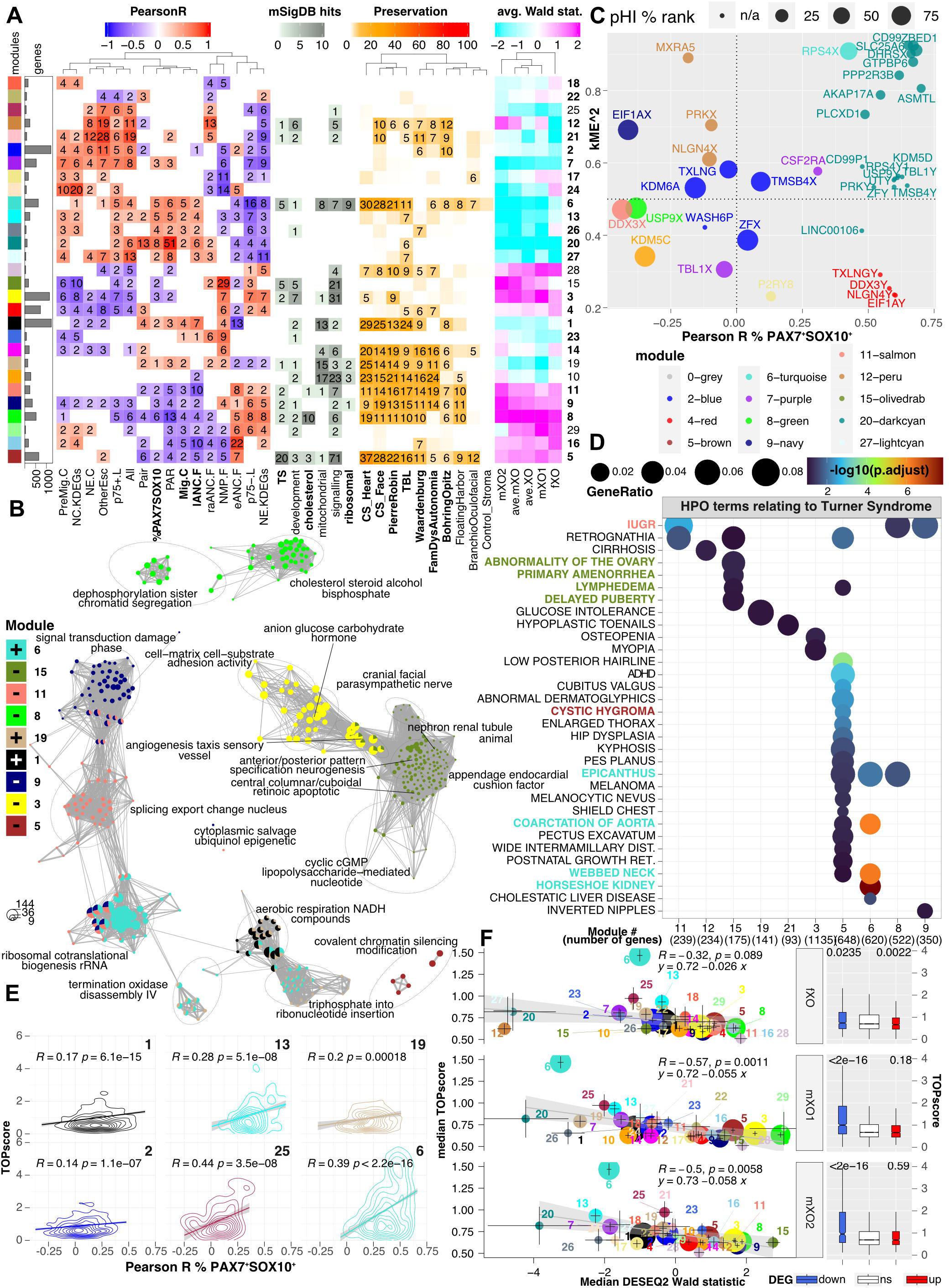
Monosomy-X sensitive gene modules correlate in human development and hiPSC-derived NCC models of monogenic neurocristopathy. **(A)** Color labels and genes per WGCNA module (1-29) with corresponding: **(i)** Correlation matrix (Pearson R) of module eigengene to karyotype status (euploid, 45,X) and %PAX7^+^SOX10^+^, as well as averaged marker set expression, and sex-linked genes (PAR, X/Y-pair, other escapees, and “All” classes combined). Only for significant correlations (p≤0.05, Fisher transformation) shown, with log-scaled adjusted p-values to the nearest integer. **(ii)** Significantly**-**enriched (p.adjust≤0.1, hypergeometric distribution) gene terms from the Hallmark and canonical pathways collections (MSigDB) representing metabolism and development. **(iii)** Preservation statistics (integer Z-score) in transcriptomes of the developing human face ^26,27^ (Carnegie stages CS13-17 & 22) and heart^25^ (CS13-23), alongside hiPSC-derived NCC models of monogenic neurocristopathy syndromes (Pierre-Robin/*SOX9* ^29^, Waardenburg/*SOX10* ^28^, Familial Dysautonomia/*IKBKAP* ^30^, Bohring-Opitz/*ASXL1* ^31^, Floating-Harbor/*SRCAP* ^32^, and Branchio-Oculofacial/*TFAP2A* ^33^). Monosomy-X human iPSC-derived trophoblast-like cells (TBL ^12^) and pancreatic tumor stroma (Stroma ^60^) as respective positive and negative controls. **(iv)** Module-averaged Wald statistic. **(B)** Enrichment map of significantly enriched gene ontology terms (GO, p.adjust≤0.01, hypergeometric distribution) as nodes. Genes shared between nodes (≥10%) depicted, based on semantic pairwise-term similarity (Jaccard distance) to cluster nodes and summarize labels. Nodes colored as pie charts by fraction of enriched modules under that term. **(C)** Module eigengene correlations (kME^2) for each sex-linked gene over its correlation with the PAX7^+^SOX10^+^ rate (colored by assigned module, sized by %pHI rank). **(D)** Dotplot of significantly-enriched (p.adjust ≤0.1, hypergeometric distribution) TS-HPO terms (x-axis, numeric module labels, above gene total). Dot color and size denote log-scaled adjusted p-value and fraction of term-associated genes per module. **(E)** Modules with significant correlation (R, with regression line and gray 95% C.I.) between genes’ TOPscores (y-axis) and their Pearson coefficient with the PAX7^+^SOX10^+^ rate (x-axis), indicated by contour plots. **(F)** Median TOPscores over median Wald statistic by gene module (colors, size relative to gene total) with standard error bars per module. Pearson R correlation (black line, 95% C.I.) and p-value indicated for each comparison (fXO, mXO1 & mXO2), alongside TOPscores by DEG category (right).

If the metabolic processes reflected in these modules play important roles in neural crest development, we would expect them to be detectable in tissues populated by the neural crest. We tested this hypothesis in transcriptomes collected from the developing heart ^25^ and face ^26,27^, spanning Carnegie stages 13-23. Both tissues (‘CS_Heart’ with 11/28 preserved modules, and ‘CS_Face’ with 12/28) show moderate to high module preservation (Z-score ≥ 5), 8 of which also co/anti-correlate significantly with the PAX7^+^SOX10^+^ rate in our hiPSC-derived NCCs (Fig. 4A). Likewise, we assessed module preservation in hiPSC-derived NCCs of monogenic conditions considered neurocristopathies: 15 modules were preserved in Waardenburg ^28^ (*SOX10*), 12 in Pierre-Robin^29^ (*SOX9*), and 11 each in Familial Dysautonomia^30^ (*IKBKAP*) and Bohring-Opitz^31^ (*ASXL1*) syndromes, most of which also overlapped with each other. In contrast, only respectively 6 and 4 modules were preserved in hiPSC-derived NCC models of Floating-Harbor^32^ (*SRCAP*) and Branchio-Oculofacial^33^ (*TFAP2A*) syndromes, while no module met the Z-score threshold in a pancreatic tumor stroma control dataset. In sum, our analysis indicates that monosomy-X sensitive modules that reflect metabolic disturbance also co-vary in relevant tissues with developmental time and in NCC *in vitro* models of canonical neurocristopathies.

Interestingly, three groupA modules (13,20,27) were exclusively preserved in our hiPSC-derived monosomy-X trophoblast model ^12^, and correlated with escapee levels (PAR, Pair & ‘otherEsc’), as well as each other (Fig. 4A, S4C), raising the question in which modules these genes were over-represented. Indeed, module 20 was strongly enriched in PAR genes (Fig. S4C) but lacked any other significant MSigDB or GO annotations (Table S2). Likewise, most gametolog pairs were dispersed across modules 4 & 20, or remained unassigned. One notable exception was the only ribosomal protein to be encoded by two distinct genes, as *RPS4X* was assigned to the translation-enriched module 6, which appeared generally down-regulated in X-monosomic lines (Fig. 3B, Fig. 4A, Table S1).

To prioritize candidate dosage-sensitive genes shared by X and Y, we squared their correlation with the averaged expression of their respective modules and plotted this coefficient (kME^2) over their individual correlation with the PAX7^+^SOX10^+^ rate (Fig. 4C). We also present their percentile-ranked probability of haploinsufficiency (pHI % rank, from^34^) as a constraint metric to reflect possible selection. Most of the highly pHI-ranked genes correlate poorly with the PAX7^+^SOX10^+^ percentage, in contrast to most PAR genes, as well as Y-linked gametologs, that correlate strongly but rank expectedly low by pHI ^35^, by the residual variation intolerance score (RVIS ^36^), or remain unranked (Y-linked gametologs). Yet, *ZBED1, SLC25A6, RPS4X* and *PLCXD1,* rank in the top half of pHI or RVIS scores and correlate strongly with the PAX7^+^SOX10^+^ rate and their kME (Fig. 4C). While homozygous loss-of-function mutations have revealed *PLCXD1* as dispensable in humans ^37^, *ZBED1* was also a top candidate in our 45,X hiPSC-derived trophoblast analysis ^12^. *RPS4X* was especially notable given that translation was over-represented in its assigned module 6, which was also enriched in the three highest-scoring HPO terms associated with TS: ‘Horseshoe kidney’, ‘Coarctation of the aorta’ and ‘Webbed neck’ (Fig. 4D). Altogether 32/114 (28.1%) of all TS-associated HPO terms were significantly enriched in 10 modules (p.adjust≤0.1, hypergeometric distribution) (Fig. 4D, S4D), and 9 of these 10 modules were also preserved in monogenic neurocristopathies (Fig. 4A).

Given the strong enrichment of ribosomal protein (RP) transcripts in module 6, and their representation in the TS-associated HPO terms (Fig. 4D, Table S2), we sought to reconcile their uniformly dampened expression with activation of mTOR signaling in 45,X NCCs (Fig. S4B). Active mTOR boosts ribosome biogenesis by relieving the repressive impact of LARP1 on translation of mRNAs with 5’ UTR terminal oligopyrimidine (TOP) motifs ^38^. However, LARP1 also stabilizes TOP-motif transcripts via the small 40S ribosomal subunit, and depletion of either LARP1 or 40S subunit RPs consequently destabilizes TOP-motif transcripts ^39^. Because *RPS4X* is an essential member of the 40S subunit, and is significantly reduced alongside *RPS4Y1* in male-derived 45,X NCCs (Fig. S2G), we asked whether TOP motif scores (“TOPscores”, from ^40^) were specifically predictive in our WGCNA or differential expression analysis. We find module 6 transcripts to feature significantly stronger TOPscores than the unassigned module, or indeed any of the other modules, even when removing RP genes (Fig. S4E, Mann-Whitney U p = 2×10^-56^ and p = 1.7×10^-32^ without RP genes). Many non-RP transcripts also contain TOP motifs, including those coding for mitochondrial proteins ^41^. Indeed, several modules show a significant correlation between their transcripts’ TOPscores and the PAX7^+^SOX10^+^ rate (Fig. 4E, S4E), including modules 1, 6, 19 and 25, which are also significantly enriched in nuclear-encoded mitochondrial proteins (Table S2). To determine whether TOP transcripts were specifically down-regulated in 45,X NCCs, we therefore compared the TOPscore of DEGs (down, up) to non-significant (ns) genes. Remarkably, median TOPscores across all WGCNA modules correlate significantly with the Wald statistic (Fig. 4F), and TOPscores of down-regulated transcripts are significantly higher (Mann-Whitney U p=0.0235, p<2×10^-16^ for respective fXO and mXO1/2) than those of unchanged or up-regulated transcripts. In sum, these data support a specific depletion of TOP motif transcripts in 45,X NCCs, and are also consistent with an essential role of mTOR signaling in NCC specification, as recently reported ^42^.

## DISCUSSION

Despite the relatively small number of genes shared by extant mammalian X & Y, to-date only TS-associated short stature has been linked conclusively to haploinsufficiency of one such gene, namely pseudoautosomal *SHOX* ^43^. This example also highlights challenges in modeling TS in rodents, as murine *SHOX* ortholog *Shox2* is autosomal ^44^ and thus remains unaffected in 39,X mice, a pattern shared with all but two murine orthologs of human PAR1 genes ^9^. Likewise, more pan-mammalian gametologs pairs were retained on both X&Y in humans than mice ^3^, with the remaining murine gametologs appearing largely haplo-sufficient in mouse cardiovascular development ^10,11^.

Given the craniofacial and cardiovascular phenotypes in TS, new models are thus needed to understand how monosomy-X impacts development, specifically in cells and tissues derived from the neural crest. Our study applies a well-established hiPSC to NCC differentiation model ^17,18^ to demonstrate that monosomy-X significantly reduces NCC specification relative to euploid controls (Fig. 1, S1), which is also reflected in expression of various NC marker sets across three independent isogenic panels (Fig. 2, S2). Differential expression indicates monosomy-X alters important metabolic pathways, specifically increasing cholesterol biosynthesis genes, while reducing RP and nuclear-encoded mitochondrial transcripts (Fig. 3, S3, Table S1). This finding is consistent with recent reports revealing metabolic reprogramming to play a key role in transitioning NCCs from proliferative to migratory and terminally differentiated states^45,46^. During the epithelial-to-mesenchymal transition (EMT), NCCs increase both ribosome biosynthesis ^45^ and aerobic glycolysis ^46^. While glycolytic genes were upregulated in our 45,X NCC transcriptomes, down-regulated transcripts included genes in oxidative phosphorylation, which has also been linked to NCC specification ^47^ (Fig. 3, Table S1). In addition, cholesterol biosynthesis genes are significantly upregulated in 45,X NCCs, which while critical for modulating NCC signaling ^48^ may also alter their patterning ^49^, and is independently consistent with hypercholesteremia seen in TS patients ^5,24^.

This link to TS phenotypes is also reflected in TS-associated HPO terms that were enriched in GSEA (28/114) and WGCNA (32/114) results (Fig. 3,4). Indeed, three TS-associated HPO terms were most significantly over-represented in module 6, which comprised transcripts for mitochondrial and ribosomal proteins, as well as other cytosolic translation factors that appeared to be reduced in 45,X NCCs. Because these transcripts feature strong TOP motifs, which are bound and stabilized by LARP1 and the small 40S ribosomal subunit ^39^, we hypothesize that their uniform reduction interferes with NCC specification. Indeed, the TOPscore was predictive of specific modules’ correlation with the PAX7^+^SOX10^+^ rate across samples (Fig. 4E), and the likelihood that transcripts were down-regulated, rather than up-regulated or unchanged (Fig. 4F). Together, these observations are consistent with the notion that the reduction in many TOP-motif transcripts may block 45,X NCCs in a requisite step of metabolic reprogramming, despite robust mTOR activation (Fig. S3, S4, Table S1).

Interestingly, RP gene and gametolog *RPS4X* was assigned to module 6, correlated with the PAX7^+^SOX10^+^ rate, and ranks in the top half of pHI/RVIS scores among X-linked genes (Fig 4). Although *RPS4X* is not implicated in cardinal TS phenotypes like short stature and gonadal dysgenesis ^50,51^, we speculate that lower overall *RPS4X* and *RPS4Y1* dosage in 45,X NNCs may reduce the production of 40S subunits, and thereby destabilize LARP1-bound TOP transcripts required in NCC specification. In support of this notion, male-derived 45,X NCCs also reduced *RPS4X* alongside other RP genes (Fig. S2G). One potential outcome of impaired ribosome biogenesis is p53 activation, as in Diamond-Blackfan anemia due to haploinsufficient autosomal RP genes, or craniofacial Treacher-Collins syndrome (*TCOF1*) due to defective rRNA production^52^. Indeed, the p53 pathway is significantly upregulated in our male-derived 45,X NCCs (Fig. S3, Table S1) and enriched in module 3 (Fig. S4B, Table S2). In contrast, female-derived 45,X NCCs maintained a high degree of *RPS4X* expression from their single X, were less impacted overall (Fig. 1), and did not upregulate the p53 pathway (Fig. S3), which may point to variation in *RPS4X* levels as a potential source of previously noted variability of NC-related TS phenotypes ^5^.

To-date, escape of *RPS4X* from XCI has only been reported for primates ^53^, which have also maintained *RPS4Y* on the Y ^3^. Our study therefore raises the question whether neural-crest related TS phenotypes may be more penetrant in mammals that also maintained Y-linked *RPS4Y*, which may be resolved by comparative cardiovascular studies of mammals with sufficiently viable monosomy-X that have a largely syntenic PAR but lack *RPS4Y* (e.g. horses) ^9,54–56^. It is also plausible that the neural crest may be sensitive to the dosage of multiple PAR genes and gametologs, which poses challenges to prioritizing candidate dosage-sensitive genes by mutational constraint measures that are based on the frequency of isolated loss-of-function variants in the population. The GenTAC (Genetically Triggered Thoracic Aortic Aneurysms and Other Cardiovascular Conditions) Registry reported that distal segmental Xp deletions minimally encompassing PAR1 and 21 additional genes can be sufficient for bicuspid aortic valve and aortic coarctation ^57^. Interestingly, no such left-sided lesions were observed in 13 subjects non-mosaic for 46,X,i(Xq) iso-chromosomes, pointing to increased Xq copy-number as a potential protective factor when Xp dosage is lacking. Yet, as the iXq would be subject to XCI, such protective gene(s) would also have to escape XCI. While this group of candidate protective genes is not necessarily confined to gametologs, it should be noted that it only includes two such genes (*RPS4X, RBMX2)*^58^. Looking forward, monosomy-X hiPSC-derived NCCs with otherwise isogenic euploid controls provide a tractable model to resolve long-standing questions on the dosage contributions of human PAR and gametolog pairs, and represent a platform to attribute clinically-relevant features of TS to the dosage of specific genes in neural-crest derived cell types.

## SUPPLEMENTARY FIGURE LEGENDS

**Figure S1: PAX7 and SOX10 quantification in isogenic monosomy-X and euploid control NCC differentiation panels. (A,B)** Representative immunofluorescence (IF) images of hiPSC-derived pairs of monosomy-X alongside euploid control NCCs (from male2 and female donors) show PAX7 (green), SOX10 (red), nuclei (Hoechst33342) and merged channels (100 um scale bar). **(C,D)** CellProfiler IF quantification of all images from 4-7 rounds of differentiation across all cell lines. Brackets and p-value (Mann-Whitney U test) indicate groups of cell lines compared within each isogenic panel. Panels separate PAX7 and SOX10 percentages, as well as lines from each donor. Symbols and colors denote karyotype and donor.

**Figure S2: RNA-seq samples, marker expression and allele-specific analysis for sex-linked genes. (A)** Dendrogram of RNA-seq samples segregating by donor and karyotype. **(B)** NC-relevant marker gene sets clustered by median expression across each set. Plot facetted by donor, with karyotype indicated below dendrograms. **(C)** Expression levels (vst) of *HOXA* and *HOXB* cluster genes by condition. **(D)** Vst counts of NC and neuro-ectodermal (NE) markers, distinguished in a human 3D model of folding neural tube from ^21^, were median-normalized and averaged over euploid and 45,X (EU & XO, respectively) NC samples for each (horizontal) donor panel. Significant differences in median expression denoted by Mann-Whitney P-value. **(E)** Heatmap of median-normalized vst differential for PAR, X-pair, and other escapee genes, alongside their reported escapee status ^58,61,62^. Left-most barplot annotation panels show each gene’s allelic ratio in female euploid XX lines, and the three central barplot panels the log2FoldChange for differentially expressed gene (DESeq2 p.adjust≤0.1) in the three isogenic comparisons (fXO, mXO1, and mXO2, or 0 if not significant). **(F)** Expression levels (vst) of X-linked lncRNA genes *FIRRE, FTX, JPX* and *XIST* by condition. **(G)** as in (F) but for X/Y gametolog pairs.

**Figure S3: (A)** Gene set enrichment analysis (GSEA) results for the Hallmark collection, ordered the quantile-normalized mean Wald statistic across all monosomy-X conditions (ave.mXO). X axis denotes the log-scaled GSEA adjusted p-value, and colors the normalized enrichment score of up-(red) and down-(blue) regulated genes associated with a given term (bubble size depicts the corresponding number of genes). Results shown for all individual isogenic monosomy-X comparisons (fXO, mXO1/2), averaged (ave.XO/ave.mXO) Wald rankings and representative control comparisons (fmE1/2: female-to-male euploid, and m1m2O: male1/2 X-monosomic samples to each other). **(B)** GSEA results for the KEGG collection, otherwise as in A). **(C)** GSEA for the DESCARTES single-cell atlas, otherwise as in A).

**Figure S4: WGCNA module preservation, eigengene correlation and term enrichments. (A)** Differential in Z-score preservation statistic after subtracting Z of a sample-size adjusted network (15 samples) across all karyotypes from corresponding Z of a network composed entirely of male-derived monosomy-X samples. Numeric labels and bar widths indicate module labels and size (gene total), respectively. **(B)** Dotplot of Hallmark and KEGG terms that were significantly-enriched (respectively, p.adjust≤0.05, hypergeometric distribution) in each module (x-axis, numeric module labels with gene totals in parentheses). Dot color and size denote log-scaled adjusted p-value and ratio of term-associated genes over all genes in each module. **(C)** Module eigengene correlations (kME, Pearson R) of sex-linked genes shared by X/Y or escaping XCI, alongside their reported escapee status (from ^58,61,62^), and X-chromosomal region (PAR, non-PAR). Column annotations plot significantly enriched (log-scaled p-value, Fisher exact test) classes of sex-linked and/or XCI-escaping genes by module. **(D)** Enrichment map of significantly enriched HPO terms linked to TS (p.adjust≤0.1, hypergeometric distribution) as individually labeled nodes with edges drawn for shared genes (≥2%). Nodes are colored as pie charts by modules enriched for a given term, and the fraction of genes from each module under that term. **(E)** Cumulative distribution of genes’ correlation coefficients with the PAX7^+^SOX10^+^ rate (top) by modules (colors), or genes’ TOPscores (bottom). Mann-Whitney U test p-value (p) as indicated on plot for modules with significantly lower (9,28) and higher (6,13) TOPscore than the unassigned module (0). RP genes of modules 1 and 6 plotted separately (dashed lines).

## Supplementary Tables

Table S1: Gene-set enrichment analysis (GSEA) results for all monosomy X-relevant comparisons (“condition”) across GO and selected MSigDB collections (Hallmark, canonical pathways, Human Phenotype Ontology & cell-type signatures). Lists normalized enrichment score (NES) and p.adjust for each comparison included for GSEA (ave.XO, ave.mXO, fXO, mXO1, mXO2, fmE1, fmE2 & m1m2O).

Table S2: Over-representation analysis (ORA) results for all WGCNA modules across selected GO and MSigDB collections (Hallmark, canonical pathways, Human Phenotype Ontology & cell-type signatures). Lists: ID, Description, GeneRatio, BgRatio and p.adjust for each module.

## MATERIALS AND METHODS

### Human iPSC culture

Euploid and X-monosomic hiPSC were derived from female and male donors, and characterized for pluripotency markers, as well as DNA methylation and X-chromosome inactivation in female euploid hiPSCs via continued *XIST* expression in RNA-seq as previously described ^12,63^. Human iPSCs were cultured in feeder-free conditions on GelTrex-coated plates (ThermoFisher Scientific) in mTeSR-1 media (Stem Cell Technologies) in 5% CO_2_ at 37°C. iPSCs were passaged with 0.5M EDTA at least weekly in small aggregates.

### Neural Crest Cell Differentiation

Confluent wells of iPSCs cultured on Geltrex-coated plates in mTeSR-1 medium were singularized with Accutase (ThermoFisher Scientific) and plated on Geltrex coated plates at 4.5-6 ×10^4^ cells/cm^2^ in DMEM/F-12, 1x N2 (recipe modified from Waisman Center Intellectual & Developmental Disabilities Research Center at University of Wisconsin-Madison: Insulin, 1.6 mg/mL; apo-Transferrin, 3.2 mg/mL; Putrescine, 0.5 mg/mL; Progesterone, 0.1 µg/mL; Sodium selenite, 0.16 µg/mL (all from MilliporeSigma)), 1% Glutamax, 1% Non-essential Amino Acids, 1 µM CHIR99021 (StemCell Technologies), 2 µM SB431542 (StemCell Technologies), 1 µM DMH1 (Tocris Bioscience), 20 ng/mL BMP4 (Peprotech), and 10 µM Y-27632 (Tocris Bioscience), for the first two days only. Media was changed daily until day 4 or 5 when the cells were harvested.

### Immunocytochemistry

NCCs were fixed with 4% paraformaldehyde for 10 minutes at room temperature, washed with 0.1% Triton-X, permeabilized with 1% Triton-X, blocked with 5% normal goat serum/2% BSA/0.1% Triton-X for 1 hour and incubated with mouse monoclonal PAX7 antibody (abcam, 1:200) and rabbit monoclonal SOX10 antibody (Cell Signaling Technologies, 1:200) overnight at 4°C. The cells were then washed, incubated with AlexaFluor-555 Goat-anti-Mouse and AlexFluor-647 Goat-anti-Rabbit Secondary Antibodies (Thermo Fisher Scientific, 1:500) for 1 hour at room temperature. The cells were then washed, stained with Hoecsht-33342 diluted 1:10,000, and mounted with ProLong Gold Antifade Mountant (ThermoFisher Scientific. The number of PAX7-and SOX10-positive cells was imaged with an EVOS FL Auto system (ThermoFisher Scientific).

### Cell Profiler analysis

The proportion of PAX7+ and SOX10+ cells was quantified by automated image analysis of Hoechst-, PAX7-, and SOX10-stained slides using CellProfiler version 4.2.4 ^64^. Nuclei were first identified in the Hoechst channel, then the PAX7 and SOX10 channels were used to quantify the number of PAX7+/SOX10+ double-positive, either single-positive type, and double-negative cells per image. Illumination-corrected images were used for all channels, and nuclei outside the 10^th^ to 90^th^ percentile size range for each Hoechst image were excluded. For each biological replicate, the values from 3 fields of view were averaged to determine the percentage of single-positive, and PAX7^+^SOX10^+^ double-positive cells, with an average of ∼1000 nuclei counted for each datapoint.

### RNA-seq analysis

RNA was extracted from NCCs using the PureLink RNA Mini Kit (ThermoFisher Scientific). Libraries were prepared at the UConn Center for Genome Innovation using the Illumina Strand mRNA Kit and 100bp paired-ends reads were sequenced to an average depth of 40 million reads/replicate on the NovaSeq (Illumina).

Read pairs were trimmed using fastp ^65^, aligned to the human reference genome (hg38) using hisat2 ^66^ for allele-specific analysis of phased variants using phaser ^67^, and quantified against GENCODE version (v36) using salmon ^68^. For analysis of XCI and escapee, A and B allele counts from phASER were tabulated and calls made by binomial test (lesser allele fraction, LAF > 0.1, p ≤ 0.05) for all X-linked genes.

For differential expression using DESeq2 ^69^, salmon count tables were filtered for genes with sufficient expression. Surrogate variables were estimated using the sva package ^70^, and added to the DESeq2 design testing for condition and correcting for differentiation round. Gene-set enrichment analysis (GSEA) using clusterProfiler ^71^ was performed on all genes ranked by DESeq2’s Wald statistic, as well as the average of the quantile-normalized Wald scores from the three euploid-45,X conditions (fXO, mXO1, mXO2) to ensure equal weighting.

Weighted gene co-expression network analysis (WGCNA) was performed on vst counts, using the WGCNA package ^72^, as a signed hybrid network using the biweight midcorrelation raised to a soft thresholding power of 12 (scale-free topology fit ≥ 0.85). Modules were correlated to PAX7^+^SOX10^+^ percentages and to averaged NCC lineage marker sets, which were median vst normalized to ensure equal weights across all sets. Module preservation analysis was performed against published RNA-seq datasets of the developing face ^26,27^, and heart ^25^, hiPSC-derived NCC models of Pierre-Robin ^29^, Waardenburg ^28^, Familial Dysautonomia ^30^, Bohring-Opitz ^31^, Floating-Harbor ^32^, and Branchio-Oculofacial ^33^ syndromes, as well as monosomy-X hiPSC-derived trophoblast-like cells ^12^ and pancreatic tumor stroma ^60^. Overrepresentation analysis of gene terms across WGCNA modules was performed with clusterProfiler ^71^.

## ACKNOWLEDGMENTS

We would like to thank the Cotney lab at UConn Health for early access to transcriptome data from the developing face, and Bo Reese at the UConn Center for Genome Innovation for mRNA-seq library preparation and sequencing. This work was supported by NIH grants R35GM124926 and R01HL141324 to S.F.P.

