## Supplementary figures and images for "Isogenic hiPSC models of Turner syndrome development reveal shared roles of inactive X and Y in the human cranial neural crest network"

### Figure S1

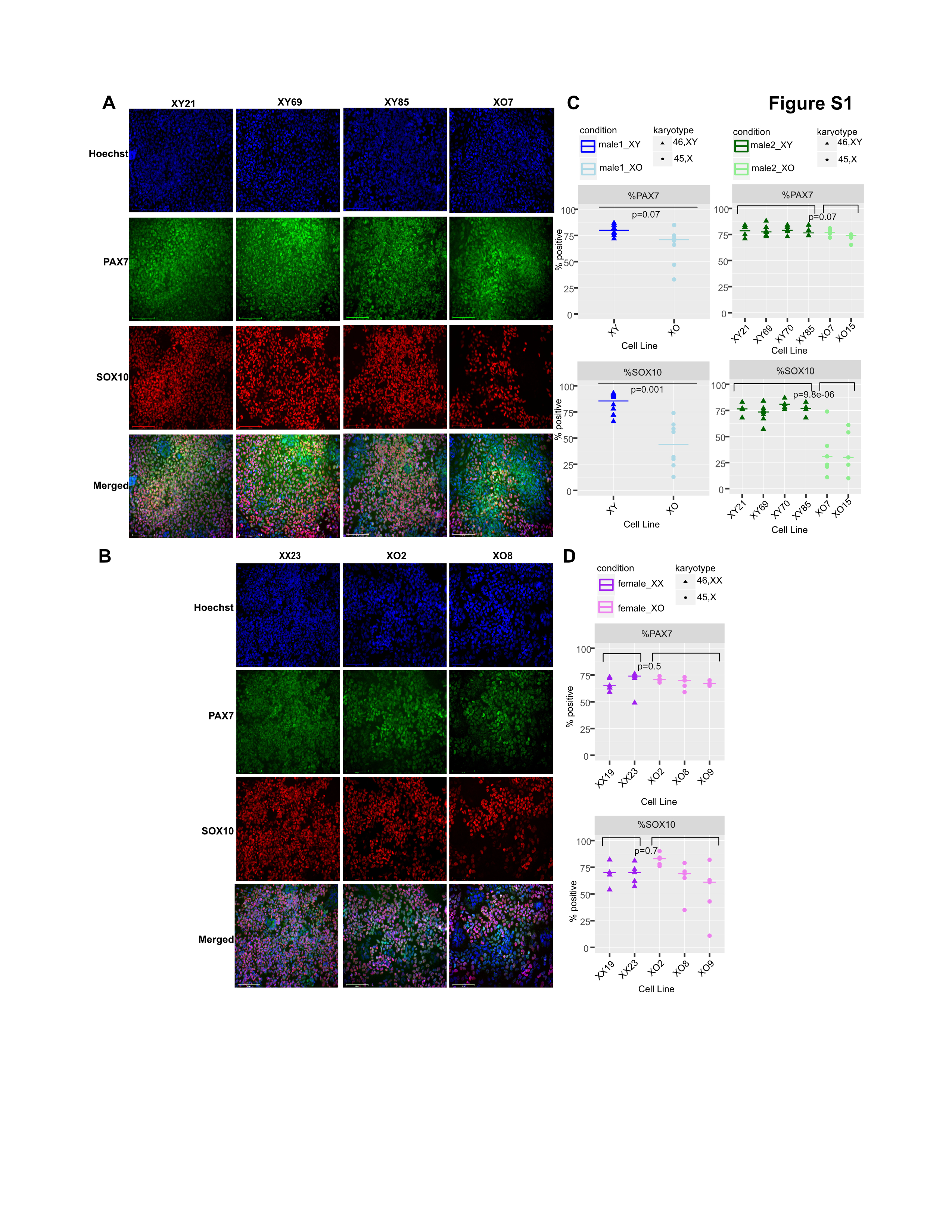

### Figure S2

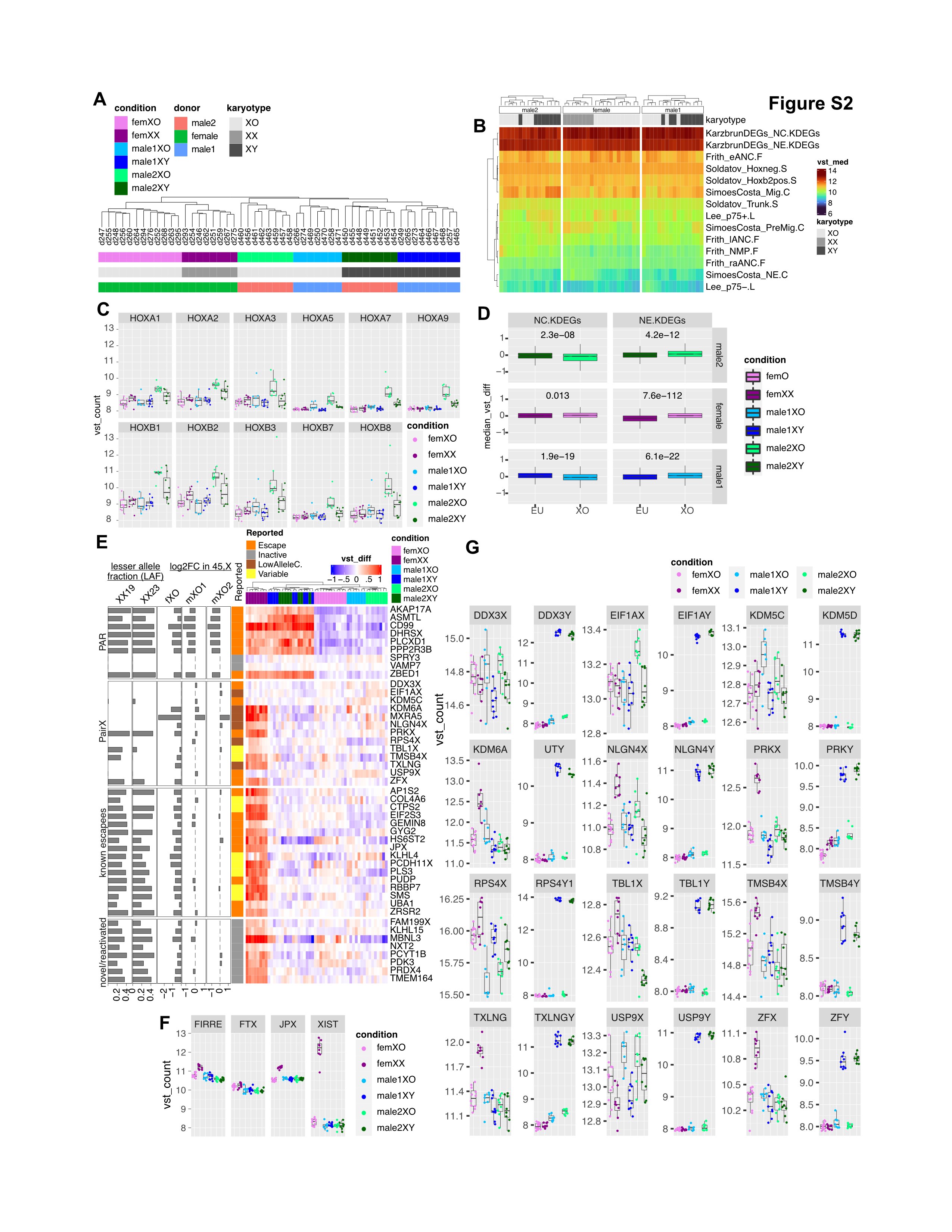

### Figure S3

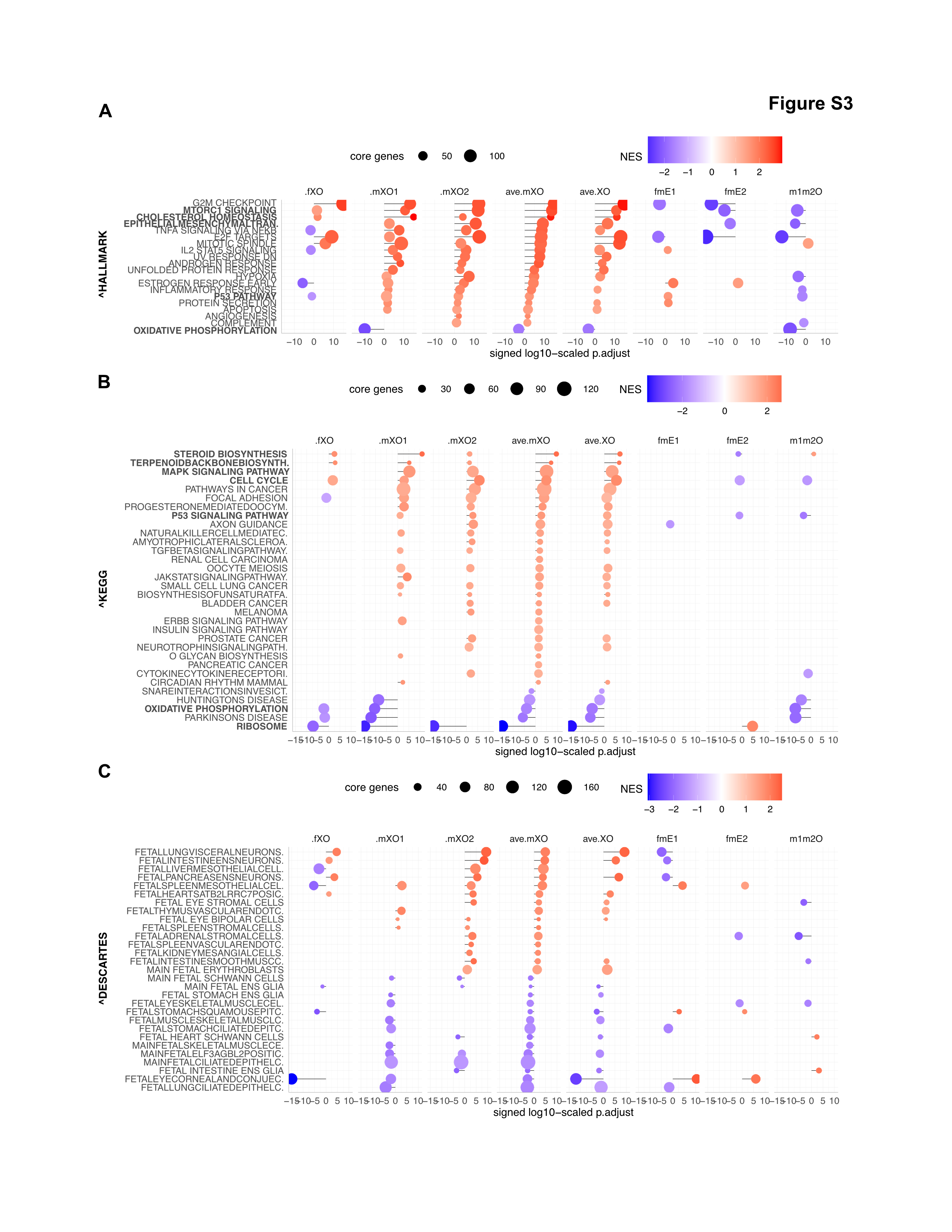

### Figure S4

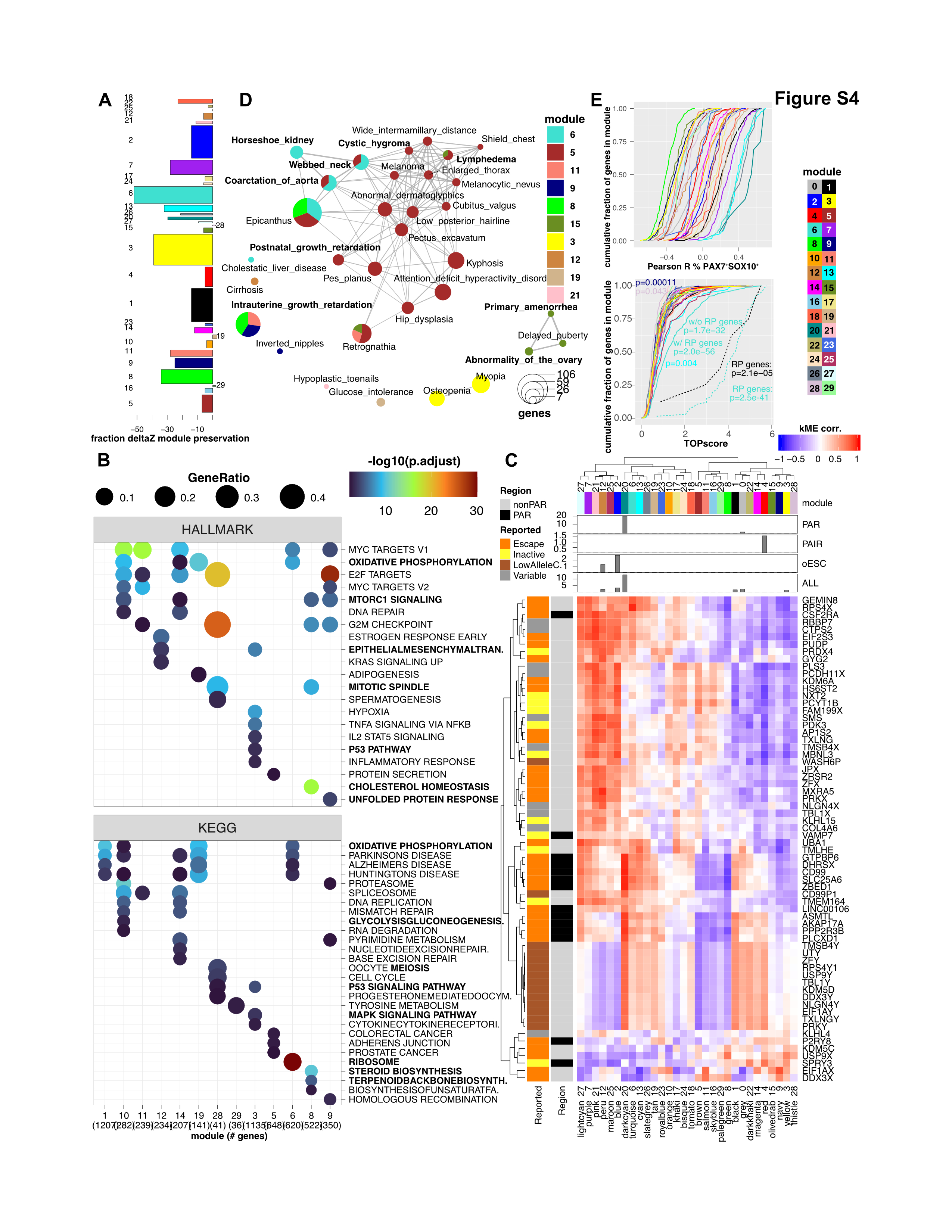
